# Phase-dependent gait robustness is not related to phase-dependent gait stability

**DOI:** 10.1101/2022.09.20.508663

**Authors:** Dinant Kistemaker, Jian Jin, Jaap H. van Dieën, Andreas Daffertshofer, Sjoerd M. Bruijn

**Affiliations:** Vrije Universiteit Amsterdam; Hohai University

**Keywords:** dynamic walker, fall prevention, walking perturbations, fall risk assessment, stability measure

## Abstract

Predicting gait robustness is paramount for targeting interventions to prevent falls. Our analysis of a compass walker model revealed that phase-dependent stability measures have limited capacity to predict overall gait robustness. Interestingly, these measures vary substantially over the gait cycle which seemingly aligns with the robustness of both humans and walking models that likewise depends on the phase at which perturbations occur. To explore this in depth, we again used a compass walker model that can walk stably and periodically and applied forward and backward perturbations (instantaneous changes in angular velocities) to the stance or swing leg at multiple instants during the single-stance phase. We then estimated the degree to which phase-dependent stability measures correlate with phase-dependent gait robustness, quantified as the maximal perturbation-induced change in mechanical energy that the walker could tolerate without falling within 50 steps. We found that these phase-dependent stability measures did not adequately predict phase-dependent gait robustness in the compass walker model. We therefore conclude that these measures are unlikely to provide reliable predictions of gait robustness or fall risk in humans.

## Introduction

Falls represent a major health threat to older adults and individuals with chronic disorders, often leading to injury, loss of independence, and increased mortality (e.g., Masud and Morris 2001; Pirker and Katzenschlager 2017). Reliable identification methods are essential to effectively target preventive interventions and should be validated through both observational and experimental studies (Bruijn et al. 2013). Several studies have estimated stability based on linearization of periodic/global (e.g., using the maximum Floquet multiplier; Hürmüzlü and Moskowitz 1987) (e.g., using the maximum Floquet multiplier; Hurmuzlu and Moskowitz, 1987) or local (e.g., local divergence rate; Ali and Menzinger 1999) behavior to evaluate stability in gait and ultimately to predict gait robustness and fall risk (Ihlen et al. 2012a; Ihlen et al. 2012b; Mahmoudian et al. 2016; Stiefenhofer et al. 2019). Here, we refer to gait *stability* as the ability of the system to return to steady-state periodic walking locomotion after an infinitesimal perturbation, whereas we refer to gait *robustness* as a measure for the maximal amount to which the system can be perturbed without falling. Floquet multipliers are the eigenvalues of the Jacobian of the stride (or Poincaré) map. This map describes how the states of a locomotor system defined on a Poincaré section (typically at the beginning of a stride) evolve from one stride to the next, (see e.g.,Wisse and Schwab 2005). The magnitudes of the Floquet multipliers indicate whether these infinitesimal perturbations grow (modulus of the multiplier >1) or decay (modulus of the multiplier <1) over successive strides and thus determine the local stability of a particular periodic gait. Apart from the challenges associated with calculating Floquet multipliers from experimental human walking data (Bruijn et al. 2013; Hurmuzlu and Basdogan 1994), the maximum Floquet multiplier does not adequately predict robustness well in several walking models (Bruijn et al. 2013). This may not be surprising, as Floquet multipliers are based on linearization and are, theoretically, valid only for infinitesimal perturbations.

In a previous modeling study (Jin et al. 2021), inspired by Norris et al. (2008), we calculated phase-dependent stability measures (e.g., local divergence rate) introduced by Ali and Menzinger (1999) from stable periodic gait cycles. These measures are derived from the local stability of the system’s limit cycle, quantified by its response to infinitesimal perturbations, and evaluated at different phases of the gait cycle. In that study, we examined whether such phase-dependent stability measures could predict overall gait robustness, operationalized as the maximum perturbation tolerated by the modeled walker (Jin et al. 2021). Specifically, we tested whether extrema of the local divergence rate or corresponding eigenvalues were correlated with the magnitude of the largest perturbation the system could withstand. The idea we tested was that, for example, a system with higher positive maximal local eigenvalue to be less. We found that these local stability measures did not correlate well with overall gait robustness. Based on these findings, we concluded that phase-dependent stability measures may not be suitable for predicting overall gait robustness.

In line with previous work (Norris et al. 2008), we found that local stability measures vary substantially over the gait cycle, even though the system remains locally unstable throughout (Jin et al. 2021). At the same time, it is well established that both humans (Cordero 2003; Eng et al. 1994; Golyski et al. 2022; Tang and Woollacott 1999) and simple walking models (de Boer et al. 2010; Williams and Martin 2020) can tolerate perturbations of different magnitudes depending on the phase at which they occur. This raises an important question: are these two phase-dependent phenomena related? For example, might phases characterized by lower maximal eigenvalues also be the phases in which the gait is more robust to external perturbations? Such a relationship would have practical implications for identifying gait phases that are particularly hazardous for fall-prone individuals. Conversely, if no relationship exists, it would further underscore important limitations of using phase-dependent stability measures to predict phase-dependent robustness, suggesting that alternative approaches may be required for assessing gait robustness and, ultimately, fall risk.

In this study, we addressed this question using a simple compass walker (Garcia et al. 1998; Norris et al. 2008). For several stable periodic gaits, we calculated the maximal local eigenvalues and trajectory-normal divergence rates across the single-stance phase. We then applied forward and backward perturbations to both the swing and stance leg at several instants during the gait cycle and quantified phase-dependent robustness as the maximal perturbation magnitude that did not lead to a fall. We then examined whether phases with smaller divergence rates or eigenvalues, indeed exhibited greater robustness.

## Methods

### Compass walker model

The compass walker model was the same as in our previous study (Jin et al. 2021) and was based on a published model (Garcia et al. 1998; Norris et al. 2008). In brief, this model consisted of two massless legs (with normalized length *l*) connecting the hip point mass *M* and foot point masses *m*, see Fig. 1. During the swing-phase, the connection of the stance leg to the ground was frictionless, effectively a hinge joint (i.e., no slip between stance foot and contact surface). Foot-strike was modelled as an instantaneous inelastic collision. The state variables were represented in Hamiltonian form as ***s*** = (*θ, φ, p*_0_, *p*_*φ*_), where *θ* denotes the angle of the stance leg with respect to the normal of the inclined floor and *φ* denotes the angle between the legs. *p*_0_ and *p*_*φ*_ are the canonical conjugate momenta:

**Figure 1.**
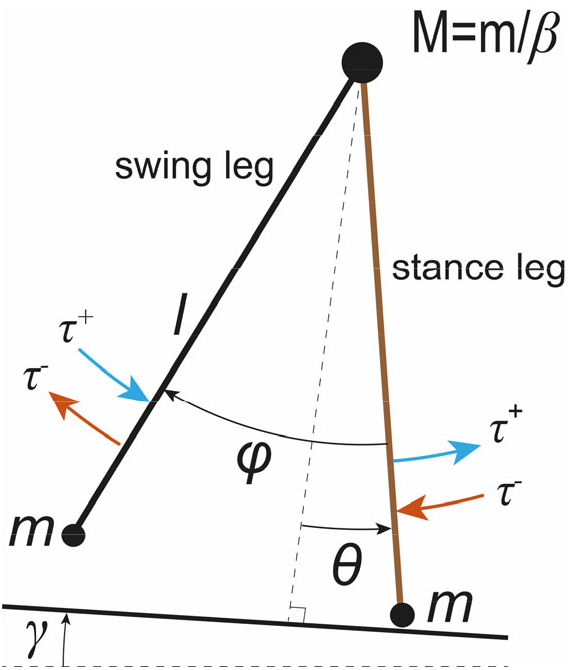
Schematic representation of the compass walker model. State variable *θ* is the angle of the stance leg with respect to the normal of the inclined plane, and *φ* is the angle between the legs. In the hip there is a point mass *M* and in the “foot” two point masses *m*. Perturbations were applied by an instantaneous forward (τ^+^) or backward (τ^+^) change in angular velocity.

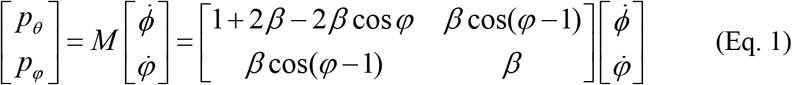

For a detailed description of the equations of motion, we refer to our previous paper [6].

The compass walker model can walk stably and periodically down a slope, with each step consisting of a single stance phase and an instantaneous double-stance phase (foot strike). Using numerical simulations (for detail see [6]), we identified a stable period-one gait for the walker model for each of the four different configurations of mass ratio 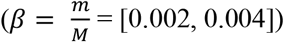 and slope (*γ* = [0.001, 0.002]).

### Phase-dependent stability measures

As in our previous study (Jin et al. 2021) and as explained in (Norris et al. 2008), we calculated phase-dependent stability measures of the swing-phase by linearizing the system in a rotating hypersurface perpendicular to the periodic trajectory. This coordinate transformation ensures that the stability estimates are invariant to perturbations along the trajectory which only produce a phase shift (i.e., a mere advancement or delay along the periodic gait cycle). From this linearization in the rotating surface, we determined the eigenvalues of the reduced Jacobian matrix (reduced because we do not consider motion in the direction along the periodic trajectory). From these eigenvalues, we computed two phase-dependent stability measures: (1) the trajectory-normal divergence rate (the sum of the eigenvalues) and (2) the maximum eigenvalue. The first measure quantifies the mean “volume” contraction or expansion rate of all infinitesimal perturbations perpendicular to the periodic trajectory (i.e., a larger trajectory-normal divergence rate indicates faster mean expansion of perturbations normal to the trajectory). The second measure quantifies the highest rate of divergence of infinitesimal perturbations along the corresponding eigenvector (i.e., a more negative eigenvalue indicates faster decay of perturbations along that eigenvector).

### Perturbations and gait robustness

For each configuration of the compass walker, and thus for each periodic gait, we applied perturbations at 50 equally spaced time instants along the periodic gait cycle. These perturbations were implemented as instantaneous forward (τ^+^) or backward (τ^-^) changes in the angular velocity (see also Fig.1) of either the stance leg 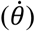 or swing leg 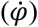. Thus, in total we investigated 4 (configurations) × 50 (perturbation instants) × 2 (perturbation directions) × 2 (legs) conditions. For each condition, we ran simulations in which we incrementally perturbed the system until the walker fell within 50 steps after the perturbation. To implement the perturbations, we first calculated 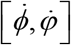 from the momenta of the unperturbed trajectory (see Eq.1):

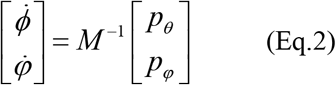

We then applied the perturbation in terms of either 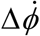 or 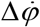, added these at the perturbation time instance to 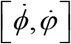 and transformed the perturbed angular velocities back to momenta using Eq.1 and carried out a forward simulation. Using this approach, we identified per leg, per perturbation, and for each of the 50 perturbation instants, the maximal perturbation (with an accuracy of 1e-4 rad√(g/l)) that could be induced without the walker falling within 50 consecutive steps.

Note that the perturbations are not forces or impulses, but step-like angular-velocity perturbations. The amount of mechanical energy delivered or dissipated during a forward or backward perturbations depends not only on the perturbation magnitude, but also on the angular velocity of the perturbed leg at the instant of the perturbation. To return to the stable periodic gait determined by the mass ratio and slope combination (Garcia et al. 1998), this change in positive or negative energy must be compensated for by the walker over subsequent strides; and, because during the single stance phase this is a conservative system, this can only happen during the collision at foot strike. Therefore, we quantified phase-dependent robustness as the change in mechanical energy induced by the maximal perturbations in angular velocity. This mechanical energy was calculated by taking the difference in the Hamiltonian (H(**s**), Eq.3; see also Norris et al., 2008), between the unperturbed and perturbed states, evaluated at the perturbation instant.

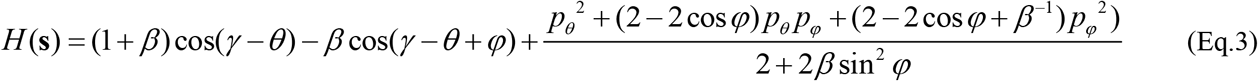

### Correlation between phase-dependent stability measures and gait robustness

We calculated correlations between both phase-dependent stability measures, i.e., the trajectory-normal divergence rate and the maximum eigenvalue, and phase-dependent gait robustness. For these stability measures to be good predictors of robustness, they do not need to show a linear relationship; any monotonic relationship is sufficient. Therefore, we used the Kendall rank correlation coefficient (Kendall 1938) rather than standard Pearson correlations. Briefly, the Kendall rank correlation coefficient (ranging from −1 to 1) evaluates the similarity in the ordering of data ranked by two variables, with values of 1 (−1) indicating perfectly monotonic increasing (decreasing) relationships and 0 indicating no association. Consistent with our previous paper, we considered Kendall rank correlation coefficients larger than 0.7 (or smaller than −0.7) to indicate strong correlations.

## Results

In Figure 2, we plotted the trajectory-normal divergence rate and the maximal eigenvalue as a function of the relative stance phase for a stable period-one gait of our compass walker with *β* set to 0.002 and *γ* to 0.001. Similar to previous findings (Jin et al. 2021; Norris et al. 2008), the divergence rate peaked at the beginning of the stance phase, drops to negative values around 25%, increases to positive values around 75%, and then rapidly declined to approximately –1 just before foot strike. The maximal eigenvalue peaked at the start of the stance phase and decreased non-monotonically to zero just before foot strike. Note that although the maximal eigenvalues varied substantially during single-stance, they remained positive throughout. This indicates that the system is locally unstable throughout the entire single-stance phase. As explained in (Norris et al. 2008), the system is nevertheless globally stable due to the inelastic collision at foot strike.

**Fig. 2.**
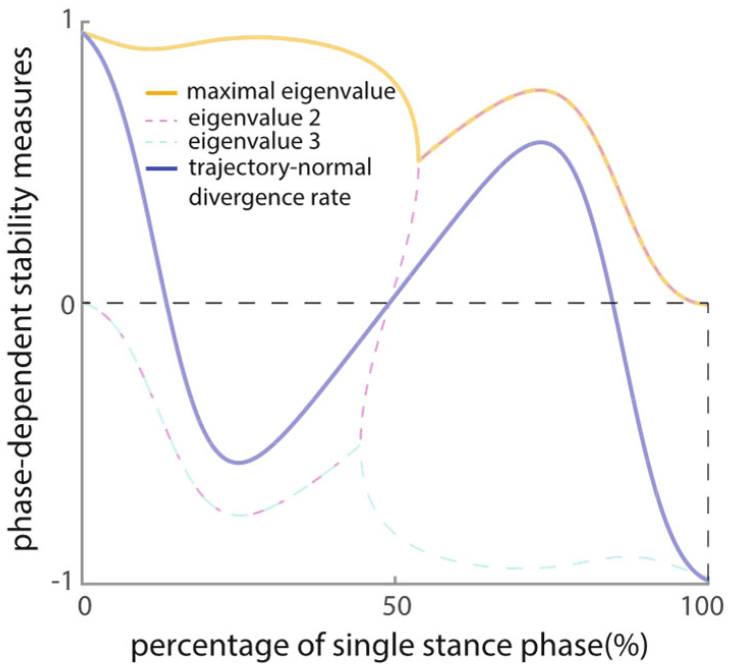
Maximal eigenvalue (yellow curve) and trajectory-normal divergence rate (blue solid curve) over the single stance phase for a walker with parameters β=0.002 and γ=0.001.

In Fig.3, we show the model’s robustness to perturbations (in terms of maximal absolute change in energy) applied to the stance and swing legs for *β* = 0.002 and *γ* = 0.001. The figure demonstrates that robustness depends on the phase especially for the swing leg. For stance-leg perturbations, maximal robustness occurred around mid-stance. Also, the walker tolerated substantially larger forward than backward perturbations. For the swing leg, maximal robustness to forward perturbations was about four times greater than for the stance leg (note the difference in scale between Figs. 3A and 3B) and peaked shortly after the start of the stance phase, and was near zero after 40% of the stance phase. The walker hardly tolerated any backward perturbations to the swing leg, except just before collision. At that point, the maximal perturbation was four orders of magnitude larger (note the break in the y-axis) than the maximal robustness for backwards perturbations earlier in the phase (see Discussion). The results for other combinations of mass-ratio and walking slope were very similar; higher mass ratio in general increased the walker’s robustness, whereas the increase of slope had minor effects on robustness (see Appendix). For the remainder of the results we will focus only the walker with a mass-ratio *β*=0.002 walking down a slope *γ*=0.001.

**Fig. 3.**
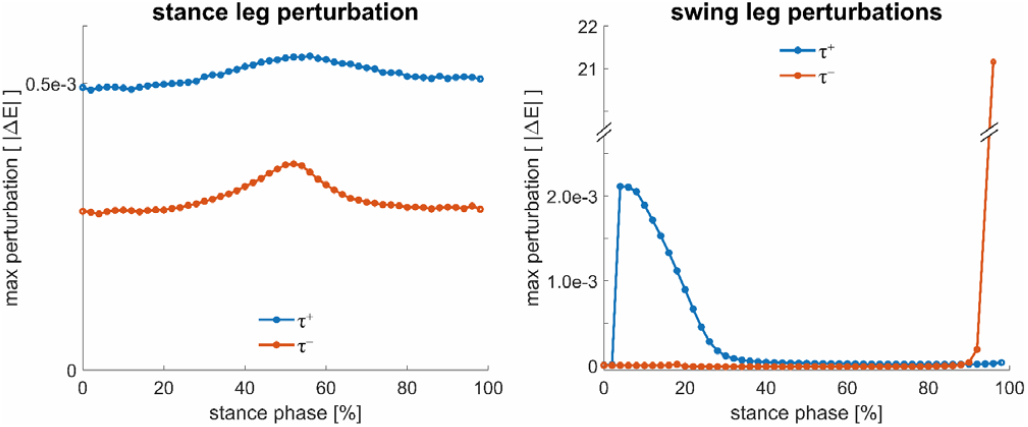
(a) Gait robustness in terms of maximal allowed absolute energy deviation for forward (blue) and backward (red) perturbations applied to the stance leg of the compass walker (for *β*=0.002 and *γ*=0.001) in different phases of the single-stance phase. (b) Same, but then for perturbations applied to the swing leg. Note that the maximal forward perturbation (just before foot strike) was four orders of magnitude greater that during other parts of the stance phase, and that of backward perturbations.

Comparing Figs. 2 and 3 suggests that phase-dependent stability measures are unlikely to serve as reliable predictors of phase-dependent gait robustness. First, they do not appear to exhibit a visible monotonic relationship with robustness.

Second, gait robustness depends not only on phase, but also substantially on the direction of the perturbation (forward vs. backward) and on which leg is perturbed (stance vs. swing). This is problematic because, at any given phase, the phase-dependent stability measures yield only a single value. In Fig. 4, we plotted gait robustness, as functions of the phase-dependent divergence rate and the maximal eigenvalue. As can be seen visually, none of the phase-dependent stability measures correlated well with phase-dependent gait robustness. This can also be appreciated from the very low Kendall rank correlation coefficients (ranging from r =0.03-0.45). Note that the somewhat higher r values for the swing leg should not be interpreted as serving a good predictor. Even though *r* was relatively high for the forward perturbation, there is essentially no useful correlation between the stability measures and robustness against backward perturbations: for most of the stance phase, robustness is almost zero, whereas the stability measures cover nearly the entire range. For the forward perturbation, stability is also not a good predictor, since the same value of a stability measure did correspond to substantially different robustness values. Because phase-dependent robustness also depended on perturbation direction, we further tested whether phase-dependent stability correlates with the sum of robustness to forward and backwards perturbations (yellow dots in Fig.4). This relationship was again found to be very weak for the stance leg. Since the robustness of the swing leg was dominated by forward perturbations (except just before collision; see Fig. 3b), we did not investigate this relationship for the swing leg.

**Fig. 4.**
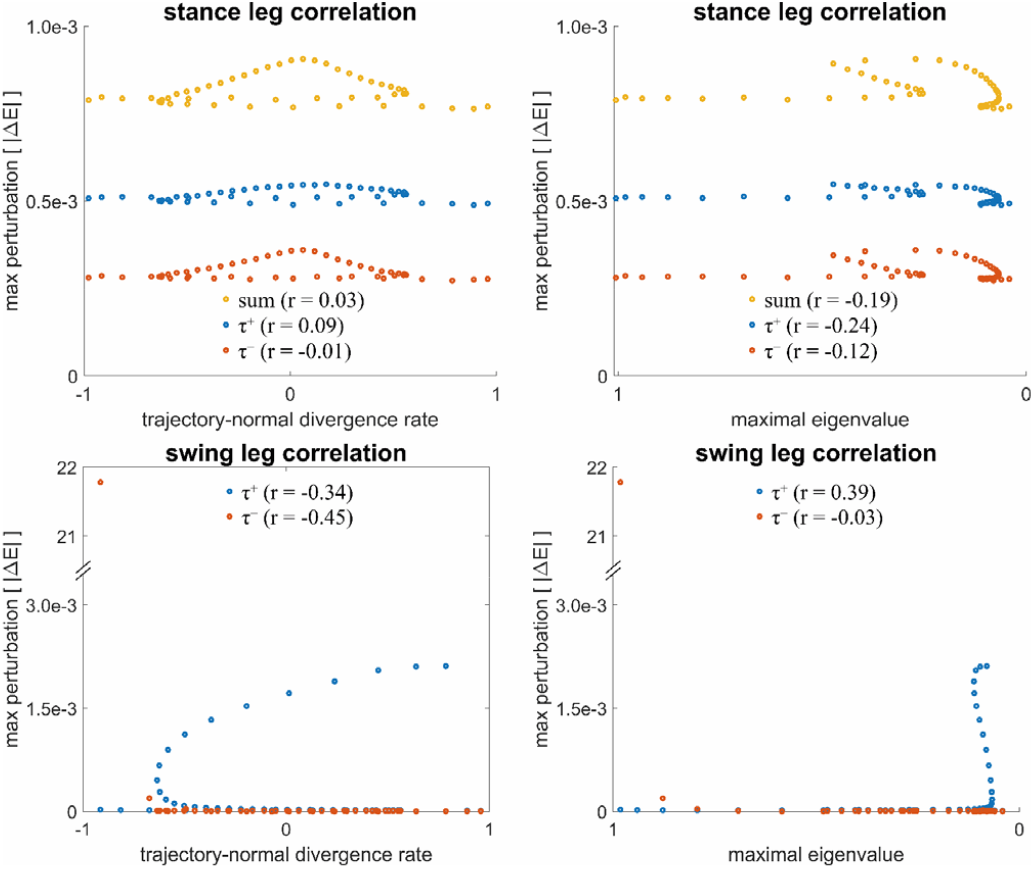
For stance leg perturbations, the relations between trajectory-normal divergence rate (a) or maximal eigenvalue (b) and maximal energy deviation for forward (τ^+^; blue) and backward (τ^-^; red) perturbations applied to the stance leg; (c)(d) For swing leg perturbations, the relations between trajectory-normal divergence rate or maximal eigenvalue with maximal energy deviation for forward and backward perturbations. Kendall rank correlation coefficients (r value) were used to quantify correlations.

## Discussion

In this study, we set out to investigate whether phase-dependent robustness, previously observed in both humans (Cordero 2003; Eng et al. 1994; Golyski et al. 2022; Tang and Woollacott 1999) and simple walking models (de Boer et al. 2010; Williams and Martin 2020), is related to phase-dependent local stability measures (trajectory-normal divergence rate and maximal eigenvalue). Studying this question in simple walker models has the advantage that the mechanics are passive and relatively straightforward. Compass walker models have proven to be valuable tools for studying fundamental principles of human walking, as demonstrated in seminal work on passive dynamic walker and simplified gait dynamics (e.g., Garcia et al. 1998; Kuo 2002; McGeer 1990). Of course, the human locomotor system is substantially more complex and walking is an active, muscle-driven, feedback-controlled process, which is not adequately captured by a passive dynamic walker. Be that as it may, our goal was not to identify predictors of human gait robustness per se, but rather to determine whether stability measures based on linearization can predict gait robustness in simple walker models. If successful, this would open a pathway for similar approaches applied to human gait. If unsuccessful, as was the case, this would underscore the difficulty of predicting gait robustness for the real human locomotor system.

During the single-stance phase, the compass walker model is a conservative system in which no energy can be dissipated; only exchange between potential and kinetic energy can occur. Consequently, the model system is marginally stable at best, and thus the maximal local eigenvalue (in the plane orthogonal to the limit cycle) must always be equal to or greater than zero (see also Norris et al. 2008). Mechanically, this means that any change in mechanical energy caused by a perturbation can only be compensated for during the foot strike collision. Thus, a trajectory that deviates from the unperturbed trajectory may still return to a steady-state gait, if a perturbation that increases mechanical energy leads to greater collision losses in the following foot strikes, or if a perturbation that decreases mechanical energy leads to reduced collision losses. This insight into the mechanics and energetics of the compass walker provides an intuitive explanation for its phase-dependent robustness. In general, the above implies that the compass walker is more robust at phases where an increase in mechanical energy results in a larger, more dissipative foot strike collision. A walker model dissipates more (or less) mechanical energy during collision when step length and/or foot strike velocity are increased (or decreased). As an example, during midstance, a forward perturbation applied to either stance or swing leg results in a larger increase in the velocity of the center-of-mass, producing larger step length and thus larger collision loss, than a backward perturbation of similar magnitude applied earlier or later in the gait cycle. In contrast, backward perturbations applied at the very beginning of the gait cycle produce step lengths that are too small, causing the walker to fall forward and resulting in low robustness to early backward perturbations. The extreme large robustness to late swing-leg backward perturbations can be explained by the fact that the induced change in angular velocity minimally affects step length (as heel strike follows almost immediately), and the additional energy is quickly dissipated by an increased collision loss.

To systematically investigate phase-dependent gait robustness and its relationship with phase-dependent stability measures, we performed simulations of periodic stable gaits of a compass walker model that was perturbed at several instants along the gait cycle. Robustness, quantified as the maximal perturbation to the stance or swing leg that did not lead to a fall within 50 steps, was found to be poorly correlated with the phase-dependent stability measures. In addition, our results clearly showed that phase-dependent gait robustness depends were substantially different for forward and backward perturbations and were also substantially difference when the perturbations were applied to the stance and swing leg, making it unlikely that phase-dependent gait robustness can be predicted from a single phase-dependent stability measure. In a previous study (Jin et al. 2021), we evaluated several stability measures (minimum, maximum, and integral of trajectory-normal divergence rate, divergence at foot strike, and Floquet multiplier) in compass-walker models to determine whether they could reliably predict overall gait robustness, assessed by a single push/pull or step-up/step-down perturbation during midstance. In line with earlier studies (Hobbelen and Wisse 2007), we found that these stability measures also did not reliably predict overall gait robustness of compass walker models.

Taken together, the present findings, combined with findings from earlier studies strongly suggest that stability measures derived from linearization are not suitable for predicting gait robustness, even in the simplest walking models. If such measures cannot reliably capture robustness in passive, low-dimensional systems with fully known mechanics, it is highly unlikely that they will succeed in the vastly more complex context of human walking, where multi-joint redundancy, muscle-driven actuation, sensory feedback, and adaptive control add substantial nonlinearities. We therefore conclude that these linearized stability measures are not likely to provide reliable predictions of gait robustness or fall risk in locomotor systems.

## Appendix A

**Fig. A1.**
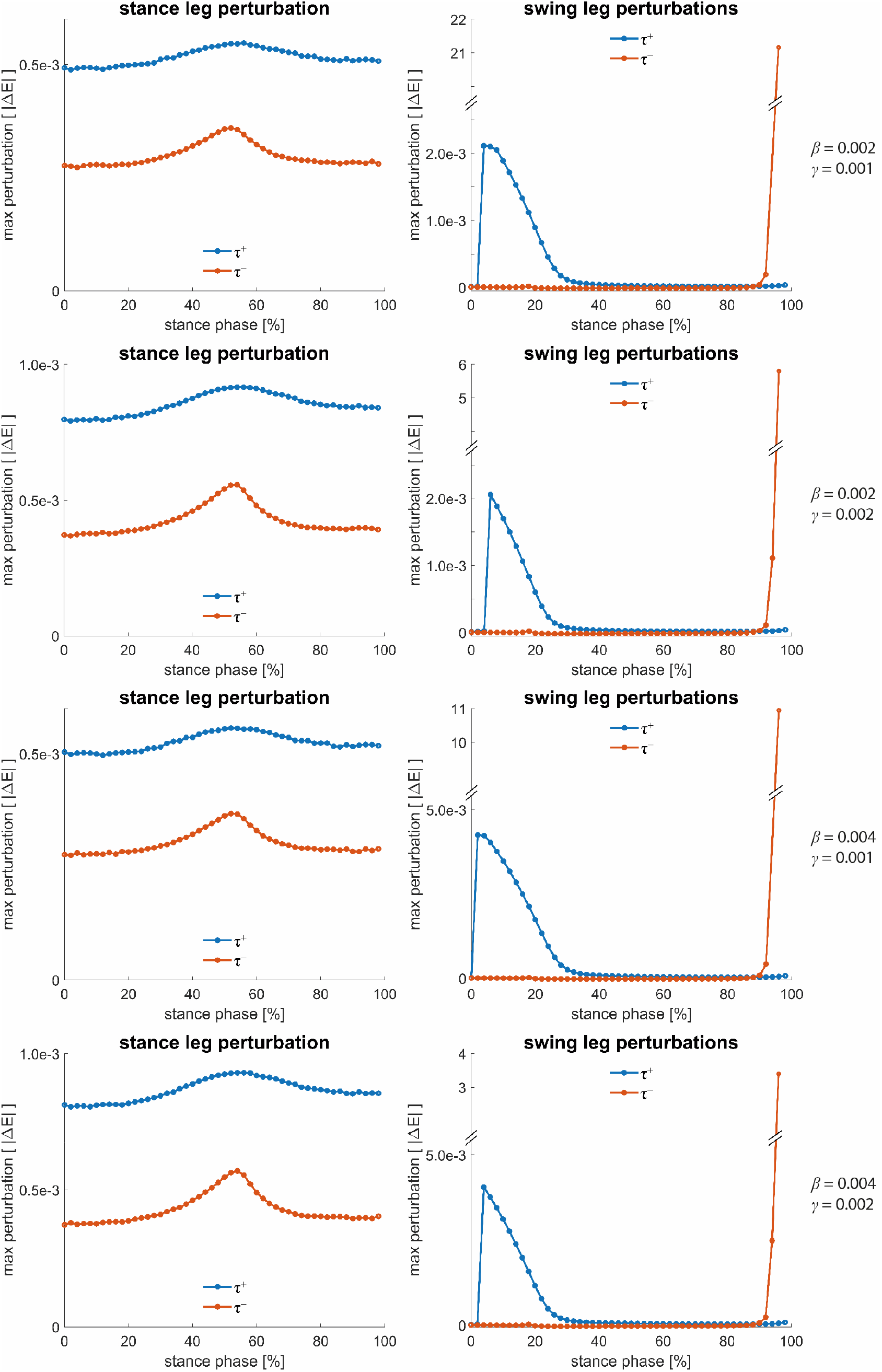
Gait robustness in terms of maximal allowed absolute energy deviation for forward (blue) and backward (red) perturbations applied to the stance and to the swing leg compass walker (for *β*=0.002&0.004 and *γ*=0.001&0.002) in different phases of the single-stance phase.

